# Identification of NanoLuciferase Substrates Transported by Human ABCB1 and ABCG2 and their Zebrafish Homologs at the Blood-Brain Barrier

**DOI:** 10.1101/2023.10.20.563277

**Authors:** Collin T. Inglut, John A. Quinlan, Robert W. Robey, Joanna R. Thomas, Joel R. Walker, Wenhui Zhou, Huang-Chiao Huang, Michael M. Gottesman

## Abstract

ATP-binding cassette (ABC) transporters expressed at the blood-brain barrier (BBB) impede delivery of therapeutic agents to the brain, including agents to treat neurodegenerative diseases and primary and metastatic brain cancers. Two transporters, P-glycoprotein (P-gp, ABCB1) and ABCG2, are highly expressed at the BBB and are responsible for the efflux of numerous clinically useful chemotherapeutic agents, including irinotecan, paclitaxel, and doxorubicin. Based on a previous mouse model, we have generated transgenic zebrafish in which expression of NanoLuciferase (NanoLuc) is controlled by the promoter of glial fibrillary acidic protein, leading to expression in zebrafish glia. To identify agents that disrupt the BBB, including inhibitors of ABCB1 and ABCG2, we identified NanoLuc substrates that are also transported by P-gp, ABCG2, and their zebrafish homologs. These substrates will elevate the amount of bioluminescent light produced in the transgenic zebrafish with BBB disruption. We transfected HEK293 cells with NanoLuc and either human ABCB1, ABCG2, or their zebrafish homologs Abcb4 or Abcg2a, respectively, and expressed at the zebrafish BBB. We evaluated the luminescence of ten NanoLuc substrates, then screened the eight brightest to determine which are most efficiently effluxed by the ABC transporters. We identified one substrate efficiently pumped out by ABCB1, two by Abcb4, six by ABCG2, and four by Abcg2a. These data will aid in the development of a transgenic zebrafish model of the BBB to identify novel BBB disruptors and should prove useful in the development of other animal models that use NanoLuc as a reporter.

**Significance Statement:** The ATP-Binding Cassette (ABC) transporters ABCB1 and ABCG2 at the blood-brain barrier (BBB) hinder pharmacological treatment of brain-related diseases. Consequently, there is a need for tools to identify BBB disruptors. We conducted a screen of ten NanoLuciferase substrates, identifying the brightest and those that were transported by human and zebrafish ABC transporters at the BBB. This work supports and complements our development of a transgenic zebrafish model, in which NanoLuciferase is expressed within glial cells, enabling detection of BBB disruption.

## Introduction

The blood-brain barrier (BBB) and its supporting neurovascular unit (NVU) regulate the delivery of circulating molecules to the human brain as well as their clearance (Daneman and Prat, 2015). Specialized endothelial cells in brain capillaries make up the BBB, and express tight junction transmembrane complexes on the apical region of the intracellular cleft, preventing the paracellular diffusion of many molecules (Chen and Liu, 2012; Pardridge, 2005). The BBB is part of a larger and more complex NVU composed of endothelial cells, astrocytes, neurons, pericytes, microglia, and the basement membrane (Banks, 2016). BBB endothelial cells express ATP-binding cassette (ABC) transporters on the luminal membrane that redirect toxins and therapeutics back into the bloodstream (Basseville et al., 2015). ABC transporters transport a wide range of substances against their concentration gradient via ATP hydrolysis (Basseville et al., 2015). Two of the highly expressed ABC transporters at the human BBB are P-glycoprotein (P-gp; also known as ABCB1) and ATP Binding Cassette Subfamily G Member 2 (ABCG2; also known as BCRP, Breast Cancer Resistance Protein). The BBB is a major hurdle when delivering therapeutics to the brain (Kadry et al., 2020). For decades, researchers have struggled to find ways to treat brain malignancies, as many therapeutics cannot successfully enter the brain and are effluxed by ABC transporters at the BBB (Durmus et al., 2015).

While the BBB and NVU are complex systems, limited drug delivery to the brain is largely attributable to ABC transporters. Deletion of the *Abcb1a* gene in mice resulted in a 20-fold increase in brain vincristine levels compared to wild-type mice (Schinkel et al., 1994). Deletion of murine *Abcb1a* and *Abcb1b, Abcg2*, or all genes at the same time has suggested cooperativity between the two transporters in keeping substrates out of the brain (Kodaira et al., 2010; Wang et al., 2018). For example, in the case of vemurafenib, a potent chemotherapeutic for cancers carrying specific BRAF mutations, deletion of *Abcb1a/b* and *Abcg2* resulted in a nearly 20-fold increase in the brain concentration compared to wild-type mice, while deletion of *Abcb1a/b* or *Abcg2* alone resulted in virtually no increase (Durmus et al., 2012). Despite the growing evidence that these transporters remain a significant impediment to the treatment of cancers and diseases of the brain, no therapies have been approved to increase the penetration of chemotherapy drugs. This may be, in part, due to limitations in model systems for the BBB. Typical *in vitro* models of the BBB are often incomplete, lacking components of the NVU, while murine and large *in vivo* models are expensive to maintain and are not amenable to high-throughput screening (Jackson et al., 2019). Thus, improved model systems to identify strategies to increase drug delivery to the brain are needed.

The zebrafish (*Danio rerio*) has emerged as a cheaper and more efficient model compared to rodents for studying the role of transporters at the BBB (Kim et al., 2017; Umans and Taylor, 2012). Similar to humans, zebrafish have a functioning BBB comprised of endothelial cells that form tight junction complexes via claudin-5 and zona occludens-1 that physically limit drug entry into the brain (Jeong et al., 2008). To block drug entry by passive diffusion through membranes, zebrafish have two homologs of human *ABCB1*—*abcb4* and *abcb5*—and four homologs of human *ABCG2*—*abcg2a, abcg2b, abcg2c* and *abcg2d* (Fischer et al., 2013; Kobayashi et al., 2008). Our lab has previously characterized the different zebrafish homologs of human ABCB1 and ABCG2 and demonstrated that Abcb4 (Robey et al., 2021) and Abcg2a (Thomas et al., 2023) have a substrate and inhibitor profile similar to those of human proteins, respectively. More importantly, we confirmed that both Abcb4 and Abcg2a are present at the zebrafish BBB (Robey et al., 2021; Thomas et al., 2023). With these functional similarities to the human BBB, and considering the potential for high throughput assays using zebrafish, they are a logical choice for a high-throughput model of the BBB (Hotz et al., 2021).

We previously developed a mouse model in which firefly luciferase, under the control of the glial fibrillary acidic protein (GFAP) promoter, was expressed exclusively in astrocytes protected by the BBB, permitting examination of ABC transporter activity (Bakhsheshian et al., 2013). Because the firefly luciferase substrate D-luciferin is transported by ABCG2, bioluminescent signal is increased only when ABCG2 is inhibited, allowing examination of ABCG2 function at the BBB (Huang et al., 2011). Recent advances in luminescence assays have resulted in the development of NanoLuciferase (NanoLuc), a more efficient alternative to traditional firefly luciferase (England et al., 2016) (Heise et al., 2013), with various substrates producing more bioluminescent light than its native substrate coelenterazine (Francis et al., 2015; Su et al., 2023; Su et al., 2020). To evaluate the feasibility of a zebrafish-based BBB integrity model with a bioluminescent read-out, we evaluated whether human and zebrafish transporters expressed at the BBB are able to transport NanoLuc substrates using a cell-based model. Previous work had shown that the NanoLuc substrate coelenterazine was transported by both ABCB1 and ABCG2 (Huang et al., 2011), but it was not known whether furimazine or other coelenterazine analogs could be transported by human or zebrafish transporters. Here, we transfect human *ABCB1* and *ABCG2*, and the zebrafish homologs *abcg4* and *abcg2a*, as well as NanoLuciferase into HEK293 cells to determine which NanoLuc substrates are transported. The development of this cell-based and potential zebrafish model could lead to the discovery of compounds that increase the permeability of the BBB via novel mechanisms.

## Materials and Methods

### Chemicals

Doxorubicin was purchased from Sigma-Aldrich (St. Louis, MO). THZ1 and Ko143 were obtained from Cayman Chemical (Ann Arbor, MI). Elacridar was purchased from Selleck chemicals (Houston, TX). Furimazine was purchased from Aobious (Gloucester, MA), ceolentrazine and coelentrazine analogues (coelenterazine-f, -h, -fcp, -hcp, -n, and -cp) were purchased from Biotium (Fremont, CA). Polyethyleneimine (PEI) was purchased from Polysciences, Inc (Warrington, PA). Flourofurimazine and Cephalofurimazine were provided by Promega (Madison, WI).

### Cell culture

HEK293 cells (ATCC, Manassas, VA) were transfected with empty pcDNA3.1 vector or with vector containing full-length *abcb4* or *abcg2a* (Genscript, Piscataway, NJ) flanked by a FLAG tag, as been previously reported (Robey et al., 2021; Thomas et al., 2023). The MDR-19 and R5 cell lines were generated from HEK293 cells transfected with full-length human *ABCB1* or *ABCG2* and have been previously characterized (Robey et al., 2008). Cells expressing the transporters were additionally transfected with a plasmid coding for NanoLuc (N1411, Promega, Madison, WI) and selected with 40 μg/ml hygromycin B (InvivoGen, San Diego, CA). Empty vector-transfected (EV) cells served as a negative control. Transfected cells were grown in EMEM medium (Mediatech, Manassas, VA) and were maintained in 2 mg/ml G418 (Mediatech) and 40 μg/ml hygromycin B. Clones expressing similar levels of NanoLuc were selected based on expression as detected by immunoblot.

### Immunoblotting

Transfected HEK293 cells were plated in a 100-mm Petri dish (Falcon) at a density of 31,250 cells/cm^2^ for 24 hours. Whole-cell lysates (20 μg) were collected in radioimmunoprecipitation assay (RIPA) buffer supplemented with 1% protease and phosphatase inhibitor and heated for 20 minutes at 37°C. Protein lysates were separated on a 4-12% precast Bis-Tris gel (NuPAGE) and transferred onto a nitrocellulose membrane. After blocking with Odyssey blocking buffer (Li-COR) for 1 hour at room temperature, blots were probed with primary antibody against FLAG (Cat F1804-200UG, Millipore-Sigma, Milwaukee, WI), ABCB1 (C219, Signet Laboratories, Dedham, MA, 1:250), ABCG2 (BXP-21, Enzo Life Sciences, Farmingdale, NY), NanoLuc (N700, Promega), or β-actin (Rabbit 4970L, Cell Signaling Technologies, Danvers, MA) overnight at 4 °C. The blot was then incubated with IRDye 680RD goat anti-mouse secondary antibody (926-68070, Li-COR, Lincoln, NE) or IRDye 800CW goat anti-rabbit secondary antibody (926-32211, Li-COR) for 1 hour at room temperature. Proteins were visualized using the Li-Cor ODYSSEY CLx (Li-COR). B-actin serviced as a loading control.

### Cytotoxicity assays

Transfected HEK293 cells were plated in opaque, white-walled 96-well plates (Corning, Corning, NY) at a density of 15,625 cells/cm^2^ and were allowed to adhere overnight. Cytotoxic drugs (doxorubicin for ABCB1 & Abcb4; THZ-1 for ABCG2 & Abcg2a) with or without ABC inhibitors (3 μM elacridar for ABCB1 & Abcb4; 5 μM Ko143 for ABCG2 & Abcg2a) were subsequently added in triplicate, and plates were incubated for 72 hours at 37 °C. HEK293 cells not expressing a transporter (Empty vector cells, EV) were used as a control. Cell viability was measured via CellTiter-Glo (Promega, Madison, WI) assay. Luminescence was measured on a microplate reader (Tecan Infinite M200 Pro, Tecan Group) and GI_50_ (50% growth inhibitory) concentration values were calculated as the concentration at which 50% luminescence was observed compared to untreated cells.

### Substrate screening

Opaque white 96-well plates were coated with polyethyleneimine (PEI, 0.025 mg/mL) for 20 minutes at room temperature to aid in cell adhesion, then washed twice with phosphate-buffered saline before cell plating. Transfected HEK293 cells were plated at a density of 31,250 cells/cm^2^. The next day, one of the ABC inhibitors (3 μM elacridar for ABCB1 & Abcb4; 5 μM Ko143 for ABCG2 & Abcg2a) or complete media was added to wells in quintuplicate and incubated for 30 minutes at 37 °C. Furimazine and its analogs, coelenterazine, or coelenterazine analogs, were subsequently added to the cells at a working solution of 470 nM for 1 minute at room temperature. Medium containing ABC transporter inhibitors and NanoLuc substrates was removed and replaced with fresh cell culture medium. After a 1-minute stabilization period, luminescence was measured on a microplate reader, every minute, for a total of 5 measurements. The brightness of the NanoLuc substrates was measured in EV cells in a similar manner, without the removal of the substrates.

### Statistics

Area under the curve (AUC) was calculated by taking the brightness values (counts per second) recorded every minute for five minutes and time (0-300 seconds at 60-second intervals) input into the trapz function in MATLAB Version 9.14.0.2337262 (R2023a) Update 5. GraphPad Prism 10 was used to determine all statistical analyses in this study, including standard deviation and statistical significance. Specific tests, number of repeats, and significance thresholds are indicated in the figure captions. Reported P values are two-tailed. One-way or two-way ANOVA statistical tests with multiple comparison tests were performed; specific tests for each assay are specified in figure captions. All experiments represent at least 3 biological replicates.

## Results

### Establishment and characterization of HEK293 cells stably expressing an ABC transporter and NanoLuc luciferase

To characterize NanoLuc substrate specificity with human ABCB1 and ABCG2, and their zebrafish homologs, Abcb4 and Abcg2a, HEK293 cells expressing these transporters were also transfected with NanoLuc. Clones were selected based on reactivity with anti-NanoLuc antibodies, as shown in **Figure 1**. We confirmed that cells co-expressing the transporters and NanoLuc still had high levels of transporter. High levels of ABCB1 and ABCG2 were expressed in cells transfected with *ABCB1* and *ABCG2*, respectively. Expression of NanoLuc was seen in the clones. Variability in the level of NanoLuc expression may be attributable to varying efficiency of transfection with NanoLuc into each transporter-expressing cell line. As we primarily aimed to compare brightness within each cell line with and without inhibitors, with minimal comparison between cell lines, this variability in NanoLuc expression level did not affect interpretation of results.

**Figure 1.**
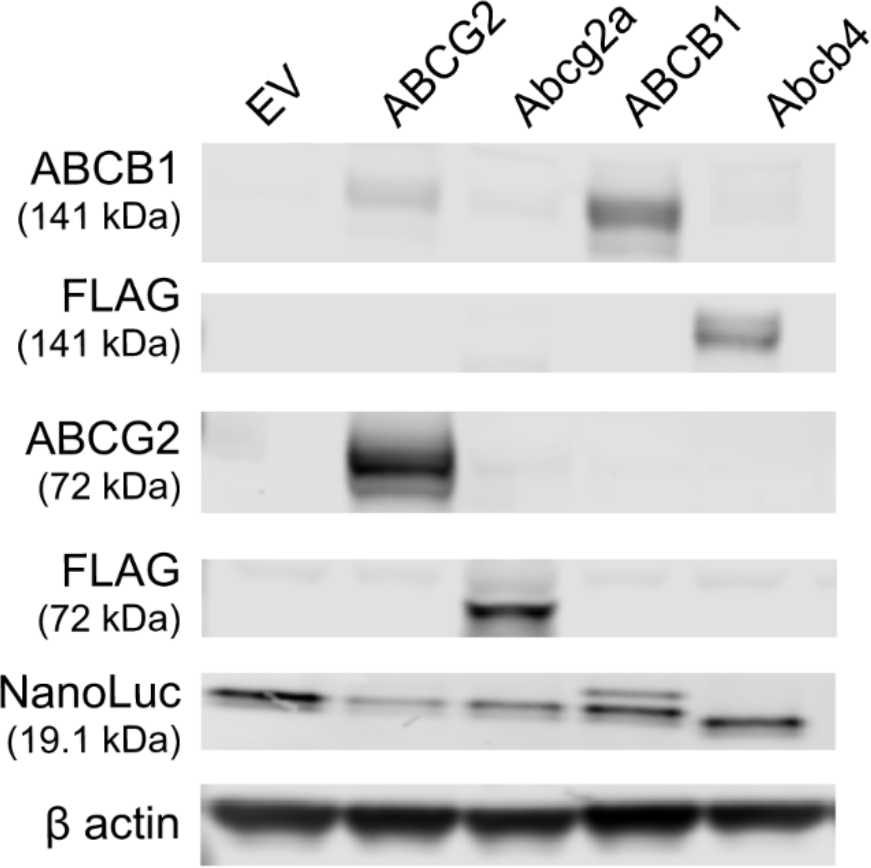
Immunoblot of four HEK293-transfected cell lines that express NanoLuc luciferase as well as human ABCG2, human ABCB1, zebrafish Abcg2a, or zebrafish Abcb4.

### ABC transporters maintain their function in double-transfected cells

To confirm the doubly transfected cell lines maintain their ABC transporter-mediated efflux capabilities, cytotoxicity assays with known substrates of ABCB1, ABCG2, Abcb4, and Abcg2a were performed (Robey et al., 2021; Sava et al., 2020; Thomas et al., 2023). Overexpression of human ABCB1 or zebrafish Abcb4 conferred resistance to doxorubicin (**Figure 2B, C**). Incubation with the ABCB1 and Abcb4 inhibitor elacridar (3 μM) sensitized both cell lines to the drug, decreasing the GI_50_ by 40.5- and 24.6-fold, respectively. Similarly, the overexpression of human ABCG2 and zebrafish Abcg2a conferred resistance to THZ1, and the ABCG2 inhibitor Ko143 (5 μM) decreased the GI_50_ by 35.6- and 16.7-fold, respectively (**Figure 2E, F**). The GI_50_ values for EV cells did not change when either drug was combined with the corresponding inhibitor (**Figure 2A, D**). Cytotoxicity results are summarized in **Tables 1 and 2**.

**Table 1.** Summary of GI_50_ values for doxorubicin in cells expressing ABCB1 or Abcb4 with or without inhibitor elacridar (3 μM) after 72 hours of treatment.

| | GI<sub>50</sub> Doxorubicin (μM) | | Fold Change |
| --- | --- | --- | --- |
|  | - Elacridar | + Elacridar |  |
| EV | 0.03 ± 0.02 | 0.03 ± 0.01 | 1.0 |
| ABCB1 | 0.81 ± 0.17 | 0.02 ± 0.01 | 40.5 |
| Abcb4 | 1.23 ± 0.42 | 0.05 ± 0.01 | 24.6 |

**Table 2.**
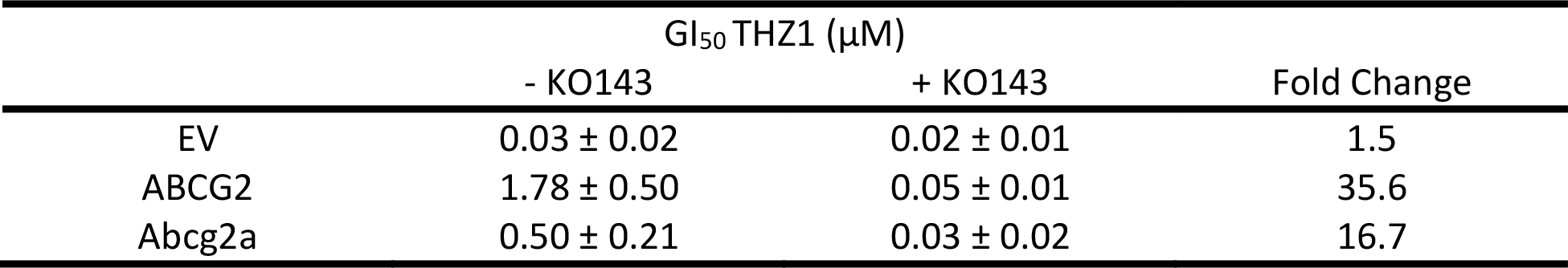
Summary of GI_50_ values for THZ1 in cells expressing ABCG2 or Abcg2a with or without inhibitor Ko143 (5 μM) after 72 hours of treatment.

| | GI<sub>50</sub> THZ1 (μM) | | Fold Change |
| --- | --- | --- | --- |
|  | - KO143 | + KO143 |  |
| EV | 0.03 ± 0.02 | 0.02 ± 0.01 | 1.5 |
| ABCG2 | 1.78 ± 0.50 | 0.05 ± 0.01 | 35.6 |
| Abcg2a | 0.50 ± 0.21 | 0.03 ± 0.02 | 16.7 |

**Figure 2.**
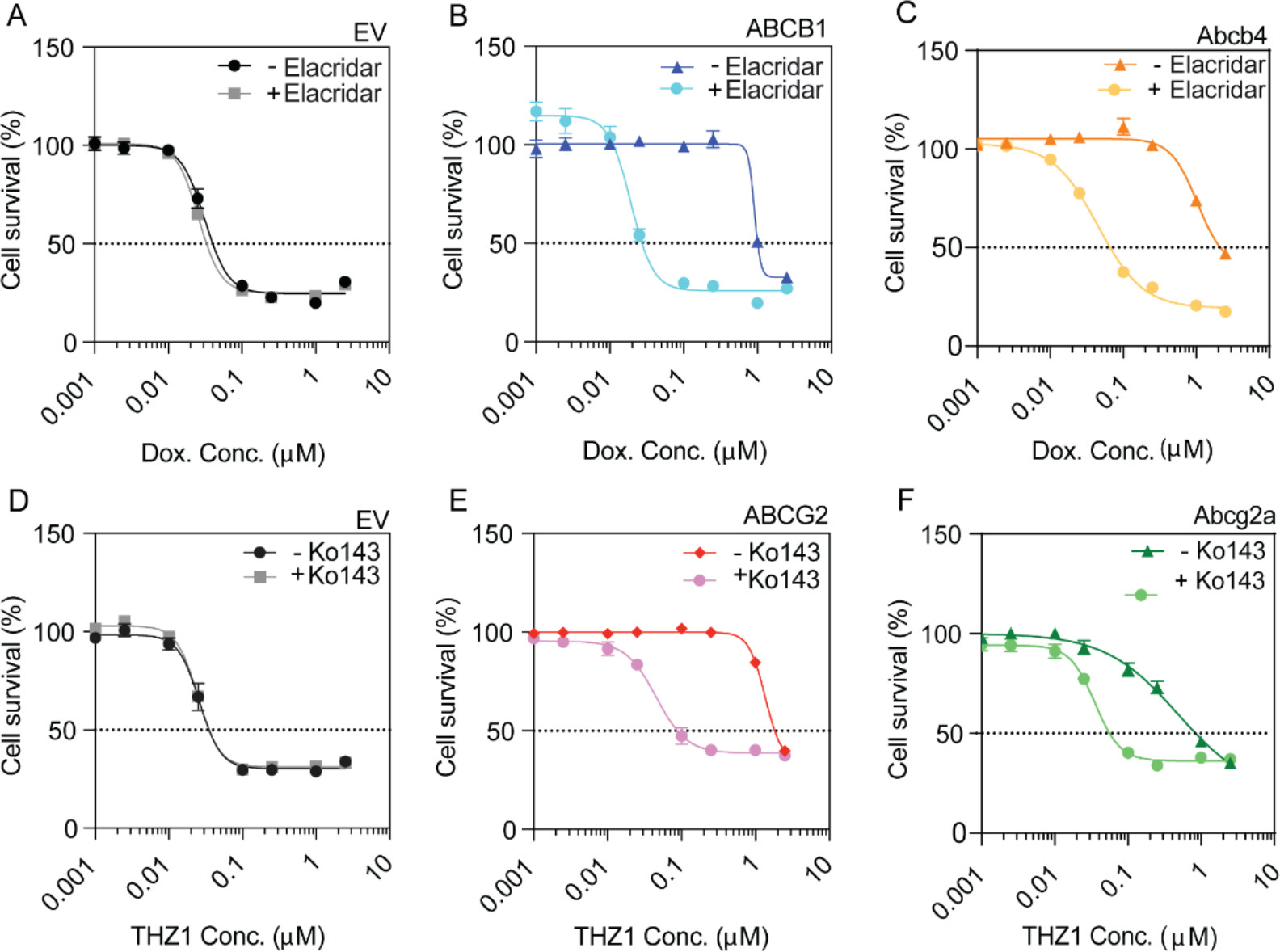
Cytotoxicity screen demonstrates doubly-transfected cell lines maintain ABC transporter function. Transfected HEK293 cells were incubated with (**A-C**) doxorubicin, an ABCB1/Abcb4 substrate, with or without the ABCB1/Abcb4 inhibitor elacridar (3 μM), or (**D-F**) THZ1, an ABCG2/Abcg2a substrate, with or without the ABCG2/Abcg2a inhibitor Ko143 (5 μM), for 72 hours.

### Brightness of Coelenterazine and Furimazine Analogs

The zebrafish larval brain at 5 days post fertilization is only 700 μm in length and 500 μm in thickness (Basnet et al., 2019; Reside et al., 2023). To identify NanoLuc substrates that may provide maximal sensitivity in the zebrafish model, we evaluated the brightness of a panel of NanoLuc substrates in EV cells. Briefly, cells were treated with the substrate of interest (470 nM) for one minute, then bioluminescence was read every minute for five minutes without removing the substrate of interest from cell culture. Area under the curve (AUC) was calculated using the resulting bioluminescence versus time data. Furimazine, the substrate specifically developed to react with NanoLuc, produced a signal 19.1 times that of native coelenterazine (**Figure 3**). Four additional coelenterazine analogs and two furimazine analogs also produced bioluminescence signal higher than that of native coelenterazine.

**Figure 3.**
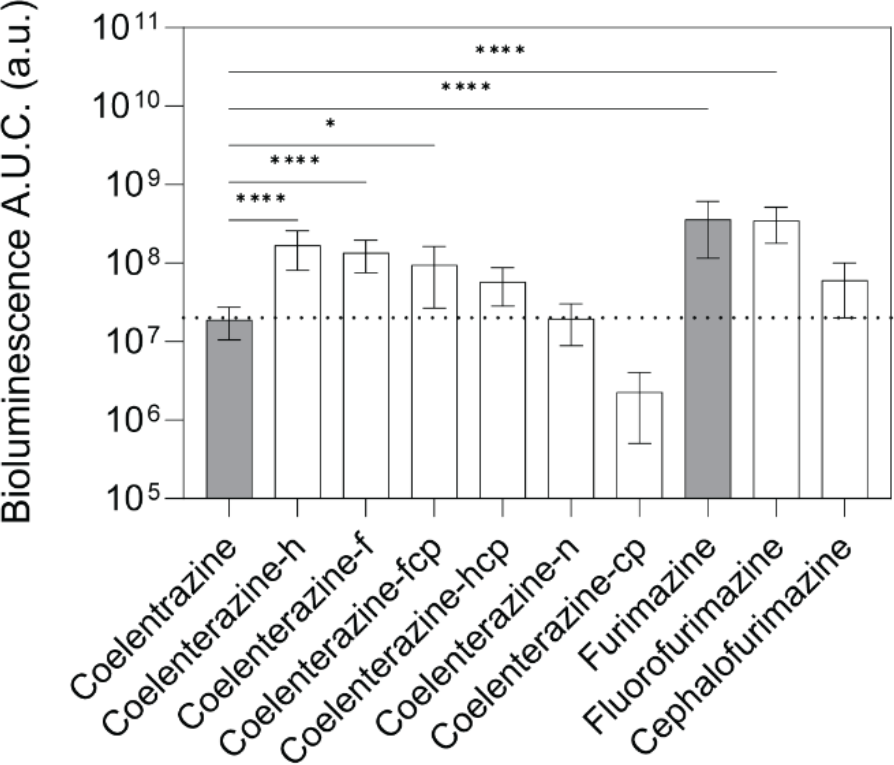
Bioluminescent signal from NanoLuc substrates in EV cells. AUC was calculated using bioluminescent signal versus time (seconds) read every minute for five minutes while cells were incubated with the NanoLuc substrate of interest. By ordinary one-way ANOVA with multiple comparisons, * indicates P<0.05, **** P<0.0001.

### ABC Transporter Substrate Status of NanoLuciferase Substrates

Next, we determined whether NanoLuc substrates are also substrates for ABC transporters, screening only NanoLuc substrates that were brighter than native coelenterazine. In addition to native coelenterazine, this included furimazine (19.1x brighter), coelenterazine-h (8.9x), coelenterazine-f (7.1x), coelenterazine-fcp (5.0x), coelenterazine-hcp (3.0x), fluorofurimazine (18.3x), and cephalofurimazine (3.2x) (**Figure 3**). Cells were incubated with inhibitor for 30 minutes (5 μM Ko143 for ABCB1 and Abcb4; 3 μM elacridar for ABCG2 and Abcg2a), then NanoLuc substrate (470 nM) for one minute before media was refreshed and bioluminescent signal was read. The bioluminescence signal was measured every minute over a 5-minute period and the area under the curve (AUC) was calculated (**Figure 4**). Native coelenterazine was previously determined to be a substrate for both human ABCG2 and ABCB1 (Huang et al., 2011) and we found that it was transported by all four ABC transporters (**Figure 4A**), as the bioluminescent signal was significantly higher in the presence of an ABC transporter inhibitor compared to the absence of inhibitor. Furimazine, which produced the highest bioluminescence signal in EV cells, was determined to be a substrate for both human ABCG2 and zebrafish Abcg2a, but not ABCB1 or Abcb4 (**Figure 4B)**. Coelenterazine-h was only a substrate for ABCG2 and Abcg2a (**Figure 4C**). Coelenterazine-f was only a substrate for ABCG2 (**Figure 4D)**. Coelenterazine-hcp was a substrate for all transporters except ABCB1 (**Figure 4E)**. Coelenterazine-fcp was a substrate of only ABCG2 (**Figure 4F**). The two furimazine analogs, flourofurimazine and cephalofurimazine, were not substrates of any transporter (**Figure 3G, H)**. The ABC transporter substrate status of each NanoLuc substrate is summarized in **Table 3**.

**Table 3.** Summary of ABC transporter substrate status of NanoLuc substrates.

| Substrate | ABCG2 | Abcg2a | P-gp | Abcb4 |
| --- | --- | --- | --- | --- |
| Native coelenterazine | * | **** | **** | * |
| Coelenterazine-h | ** | ** | - | - |
| Coelenterazine-f | **** | - | - | - |
| Coelenterazine-fcp | ** | - | - | - |
| Coelenterazine-hcp | **** | *** | - | ** |
| Furimazine | *** | *** | - | - |
| Flourofurimazine | - | - | - | - |
| Cephalofurimazine | - | - | - | - |

By ordinary two-way ANOVA with multiple comparisons, \* indicates P<0.05, \*\* P <0.01, \*\*\* P<0.001, \*\*\*\* P<0.0001. (-) indicates that the NanoLuc substrate was not an ABC transporter substrate.

**Figure 4.**
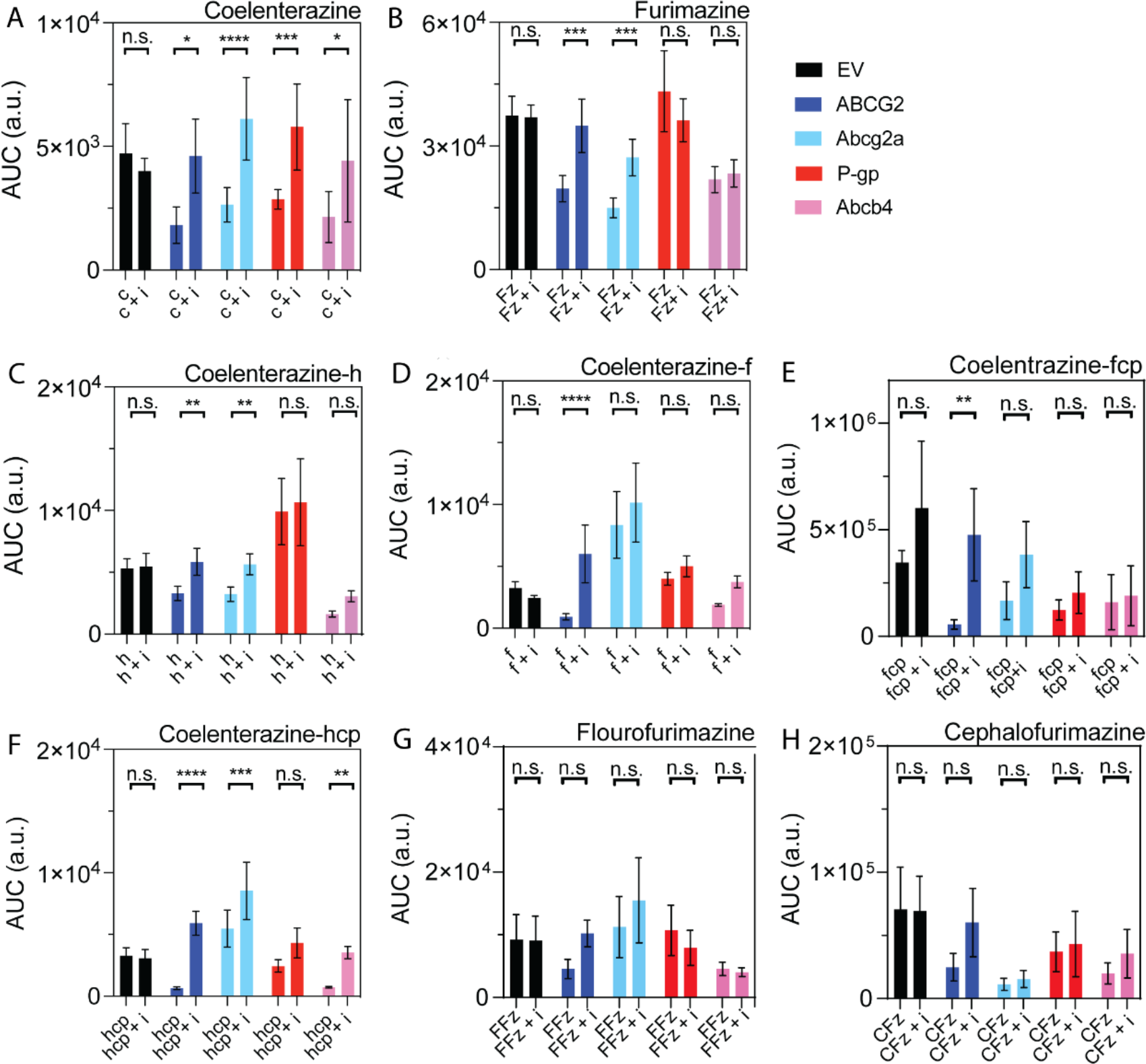
Area under the curve values from bioluminescence versus time with (+ i) and without ABC transporter inhibitors permit identification of ABC transporter substrates. HEK293 cells transfected with the transporter of interest and NanoLuciferase were incubated with inhibitor (elacridar (3 μM) for ABCB1 and Abcb4; Ko143 (5 μM) for ABCG2 and Abcg2a) for 30 minutes before introduction of the NanoLuc substrate (470 nM) for 1 minute. Bioluminescence was measured every minute for five minutes after the media were refreshed. AUC of bioluminescence versus time graphs are shown for uninhibited and inhibited (+i) (**A**) native coelenterazine (abbreviated c), (**B**) furimazine (Fz), (**C-F**) coelenterazine analogs, abbreviated by their suffix, and (**G, H**) furimazine analogs (flourofurimazine, FFz; cephalofurimazine, CFz) in a panel of transfected HEK293 cells. By ordinary two-way ANOVA with multiple comparisons, * indicates P<0.05, ** P <0.01, *** P<0.001, **** P<0.0001.

### Degree of Efflux of NanoLuc Substrates

Foreseeing potential sensitivity issues with our zebrafish model of the BBB, we also aimed to determine the degree to which various NanoLuc substrates were effluxed. A substrate that is a good NanoLuc substrate (i.e., produces a large bioluminescent signal) but a poor ABC transporter substrate (i.e., is not effluxed to a high degree) may lower specificity of the *in vivo* model. To evaluate this possibility, we looked at the luminescence ratio between cells with inhibited transporters and cells with uninhibited transporters after five minutes of efflux. By this measure, poor substrates have a low percent increase in luminescence (near 0%), while substrates that are highly effluxed will have a larger percent increase in luminescence (**Figure 5**). EV cells incubated with all the NanoLuc substrates with or without an ABC inhibitor had a ∼0% increase (**Figure 5**). The largest percent increase in bioluminescence occurred when coelenterazine-hcp was incubated with cells overexpressing ABCG2 and treated with the Ko143 inhibitor, resulting in a ∼730 ± 40.6% increase in luminescence (**Figure 5F**).

**Figure 5.**
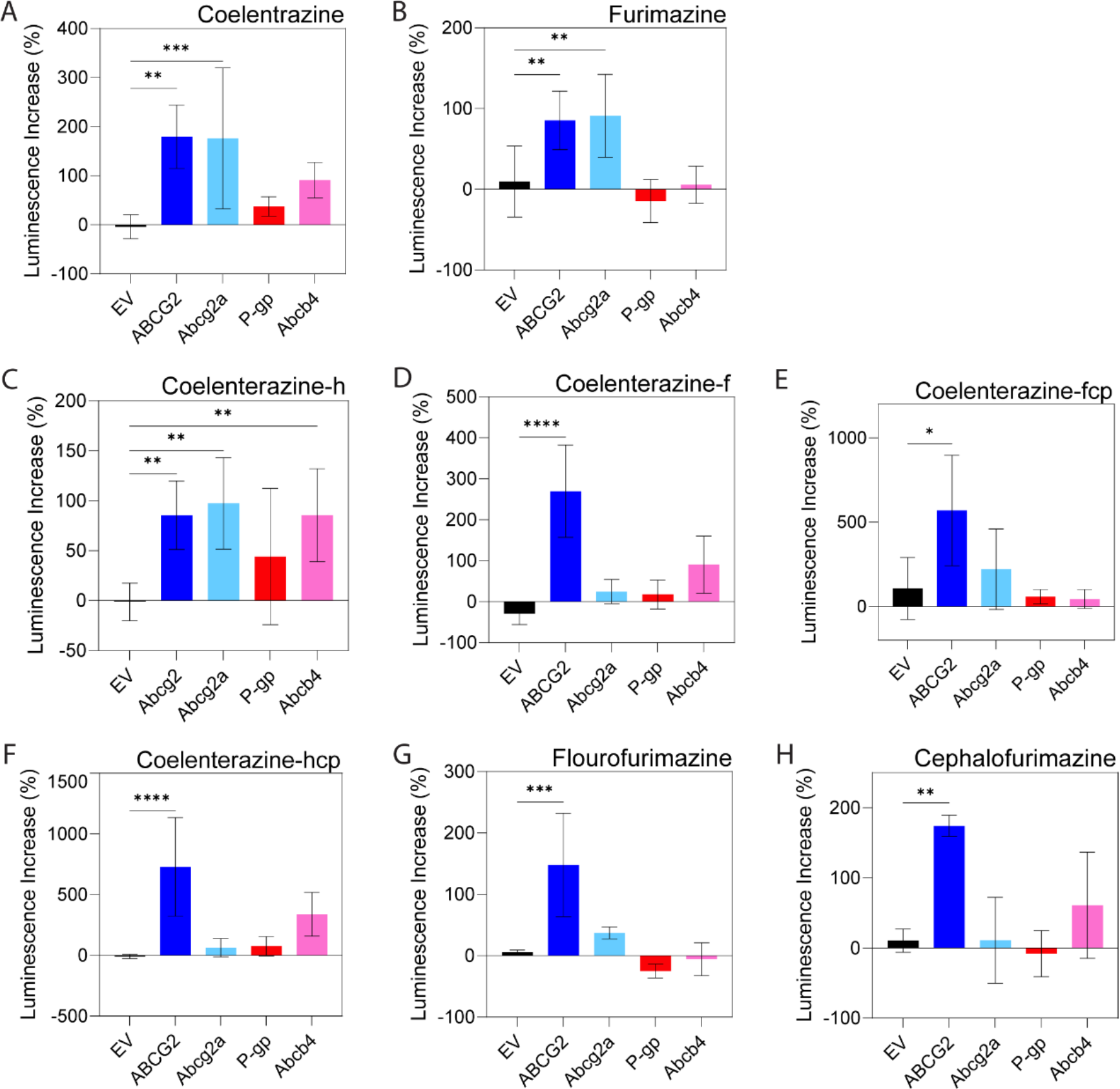
Percent increase in residual luminescence. Measurements taken after five minutes of efflux for cells treated or not treated with ABC transporter inhibitors (for ABCG2 and Abcg2a, Ko143, 5 μM; for ABCB1 and Abcb4, elacridar, 3 μM; 470 nM NanoLuc substrate of interest) for (**A**) native coelentrazine, (**B**) furimazine, (**C-F**) coelentrazine analogs, and (**G, H**) furimazine analogs. Larger luminescence increases indicate a more strongly effluxed substrate. By ordinary one-way ANOVA with multiple comparisons, * indicates P<0.05, ** P <0.01, *** P<0.001, **** P<0.0001.

## Discussion

The BBB remains a major obstacle to the management of diseases of the central nervous system (Gosselet et al., 2021), as most therapeutics cannot effectively enter the brain on their own because of the intact BBB (Bhowmik et al., 2015). BBB endothelial cell tight junctions (Luissint et al., 2012) and apical ABC transporters ABCB1 and ABCG2 prevent access to the brain (Begley, 2004) for a number of chemotherapies and targeted therapies (Anreddy et al., 2014; Choi, 2005; Shukla et al., 2011). While approaches such as focused ultrasound show promise for improving drug delivery across the BBB (Gasca-Salas et al., 2021; Rezai et al., 2020), ABC transporters may still remain a barrier to drug transport even after BBB physical integrity is compromised (de Gooijer et al., 2021; Goutal et al., 2018; Goutal et al., 2023). This challenge underscores the need to identify methodology to disrupt the BBB and ensure inactivation of ABC transporters for maximal drug delivery to the neuropil.

Our goal is to establish a zebrafish model of the BBB to screen for potential BBB disruptors in which introduction of a chemical disruptor permits entry of a NanoLuc substrate into the zebrafish brain by disrupting endothelial cells and/or inhibiting ABC transporters at the BBB. This study supports these efforts by characterizing the brightness, substrate status, and the degree to which various substrates were effluxed. We identified seven substrates brighter than native coeleterazine (coelenterazine-f, -h, -fcp, -hcp, furimazine, and cephalofurimazine) and two substrates as bright or less bright than native coelenterazine (coelenterazine-n and -cp) when reacted with NanoLuc. This informs substrate choice in the zebrafish model, enabling selection of the substrate most likely to give a strong signal to enhance sensitivity. Understanding which ABC transporter substrates are transported permits deeper probing into potential mechanisms of action of BBB disruptors, as use of specific NanoLuc substrates in the zebrafish model will elucidate the impact of BBB disruptors on specific ABC transporters. Screening experiments revealed that several NanoLuc substrates tested were also substrates for human ABCG2, excluding furimazine analogs. Only native coelenterazine was a substrate for human ABCB1. More importantly, we determined that native coelenterazine, furimazine, coelenterazine-h, and coelenterazine-hcp were substrates for zebrafish Abcg2a, while coelenterazine-f, flourofurimazine, and cephalofurimazine were not substrates for that transporter. Similarly, native coelenterazine and coelenterazine-hcp were substrates for zebrafish Abcb4. Generally, zebrafish homologs performed similarly to their human counterparts, with the only differences being transport of coelenterazine-hcp by Abcb4 and not ABCB1 and transport of coelenterazine-f and coelenterazine-fcp by ABCG2 and not Abcg2a. This varied ABC transporter substrate status profile of different NanoLuc substrates may permit probing of the role that individual transporters play at the BBB. Finally, understanding the degree to which ABC transporters efflux NanoLuc substrates may influence substrate choice. More efficiently effluxed substrates may further allow for greater specificity of the assay, supporting the use of substrates with lower brightness but different transporter status profiles.

The difference in transporter substrate status of NanoLuc substrates between human and zebrafish homologs may be in part attributable to slight differences in the amino acid residues in the ABC transporter substrate binding pocket. We previously reported that, while ABCB1 and Abcb4 have similar substrate profiles and are expressed in similar locations in humans and zebrafish, their substrate profiles are not completely overlapping. This difference in specificity is likely due to differences in the amino acids in the binding pocket of Abcb4 compared to P-gp (Robey et al., 2021), and the differences in substrate specificity found here recapitulate those results. In this study, we found differences in substrate specificity profiles between ABCG2 and Abcg2a. Our recent study indicates that Abcg2a is the zebrafish homolog most strongly correlated with ABCG2 in terms of both substrate specificity and localization of expression. We also found that ABCG2 and Abcg2a, while being functionally similar, are only 61.4% homologous in amino acid sequence (Thomas et al., 2023). While further evaluation of the similarity between Abcg2a-d and ABCG2 binding pockets is warranted, the differences in substrate specificity found here are likely attributable to slight differences in amino acid residues in the transporter binding pocket.

Native coelenterazine and its analogs contain an imidazopyrazinone core, a nitrogen-bearing heterocycle, and three side groups (Francis et al., 2015). Although there are only slight differences within the side groups corresponding to different amino acid side chains, the ABC transporter substrate status of these molecules varied greatly. Recent reports have identified two furimazine analogs capable of penetrating the BBB, flourofurimazine and cephalofurimazine, enabling enhanced imaging in the brain (Su et al., 2023; Su et al., 2020). Our findings here indicate that these compounds are not substrates of ABCB1 and ABCG2, while furimazine, a previous non-BBB penetrant iteration of the new analogs, is an ABCG2 substrate and is not BBB penetrant. This suggests that cephalofurimazine and flourofurimazine are able to penetrate the BBB because they are not substrates for the major BBB transporters ABCB1 and ABCG2.

Zebrafish offer a unique model to study transporter activity at the BBB. Our lab has recently characterized zebrafish Abcg2a and Abcb4 transporters as functional homologs of human ABCG2 and ABCB1 and both are expressed at the BBB (Robey et al., 2021). Having now established the zebrafish ABC transporter status of NanoLuc substrates, we are developing a transgenic zebrafish model that relies on NanoLuc luciferase luminescence to study the functions of these ABC transporters in determining BBB permeability. Overall, this demonstrates the feasibility of using NanoLuc substrates as a tool to probe the role of ABC transporters in BBB integrity in zebrafish. In addition, it should be noted that knowing whether various NanoLuc substrates are transported by the major multidrug transporters ABCB1 and ABCG2 will facilitate interpretation of results obtained using these new NanoLuc substrates in a variety of cell-based and whole animal assays.

## ABBREVIATIONS

ABC: ATP-Binding Cassette
ABCB1: ATP binding cassette subfamily B member 1
ABCG2: ATP binding cassette subfamily G member 2
a.u.: arbitrary units
AUC: area under the curve
BBB: blood-brain barrier
BCRP: breast cancer resistance protein
CFz: cephalofurimazine
EV: empty vector
FFz: flourofurimazine
GFAP: glial fibrillary acidic protein
i: inhibitor
NanoLuc: NanoLuciferase
NVU: neurovascular unit
PEI: polyethyleneimine
P-gp: P-glygoprotein

## Acknowledgements

We thank George Leiman and William Rhodes for editorial assistance. We thank Promega for the provision of novel furimazine analogs.

## Data Availability Statement

The authors declare that all the data supporting the findings of this study are contained within the paper.

## Authorship Contributions

*Participated in research design:* R. Robey, M. Gottesman

*Conducted experiments:* C. Inglut, J. Quinlan

*Contributed new reagents or analytical tools:* J. Walker, W. Zhou

*Performed data analysis:* C. Inglut, J. Quinlan, R. Robey, J. Thomas, H.C.-Huang, M. Gottesman

*Wrote or contributed to the writing of the manuscript:* C. Inglut, J. Quinlan, R. Robey. J. Thomas, J. Walker, W. Zhou, H.-C. Huang, M. Gottesman

All authors reviewed and agree with the final manuscript.

## Footnotes

This research was supported by the Intramural Research Program of the National Institutes of Health, the National Cancer Institute (C.T.I., J.A.Q., R.W.R., J.R.T., M.M.G.). It was also supported by the UMD-NCI Partnership for Integrative Cancer Research (C.T.I, J.A.Q., H.C.-H., R.W.R., M.M.G.)

